# L-Cystine Is a Conserved Host-Entry Signal That Activates Development in Diverse Parasitic Nematodes Through an Olfactory Receptor

**DOI:** 10.64898/2026.11.24.768902

**Authors:** Zhu Wang, Mi Cheong Cheong, Alexander H. Vidal, Manisha Mishra, George Wendt, Adler R. Dillman, Steven A. Kliewer, James J. Collins, David J. Mangelsdorf

**Author notes:** Deceased August 3, 2025 (David J. Mangelsdorf).

## Abstract

Soil-transmitted helminths infect over 1 billion people worldwide, yet how dormant infective larvae sense entry into a host and exit developmental arrest to establish infection remains largely unknown. Here, in the human parasite *Strongyloides stercoralis*, we identify L-cystine as a host signal that stimulates larval feeding and synthesis of dafachronic acid, a steroid hormone that activates DAF-12 and promotes post-infection development. A CRISPR screen of candidate receptors identified a previously uncharacterized olfactory GPCR, Ss-OR1, that is activated by L-cystine in heterologous cells and required for a normal L-cystine-induced feeding response in the parasite. Surprisingly, L-cystine also activated an OR1 homolog from the insect parasite *Steinernema carpocapsae* and stimulated feeding in its infective larvae. These findings identify L-cystine as a conserved host signal that activates OR1 homologs and stimulates feeding in nematodes infecting evolutionarily distant hosts.

**Significance statement:** Successful parasitism depends on an infective larva recognizing that it has entered a suitable host. The signals that provide this information are largely unknown. We show that L-cystine, an oxidized amino acid present in host tissues and circulating fluids, activates feeding in nematodes that parasitize either humans or insects. We also identify OR1, an olfactory GPCR responsive to L-cystine, and establish its role in the activation of *Strongyloides stercoralis* infective larvae. The conservation of L-cystine responsiveness across nematodes that parasitize evolutionarily distant hosts suggests that these parasites exploit a common chemical feature of the internal host environment. This work uncovers a previously unknown mechanism linking host entry to parasitic development and identifies a potential point of therapeutic intervention.

## Introduction

Many parasitic nematodes, including those that infect over 1 billion people worldwide, persist in the environment as non-feeding, developmentally arrested infective third-stage larvae (L3i). These larvae tolerate harsh environmental conditions but must resume feeding and development upon entering a host to establish infection. This activation is controlled by a conserved neuroendocrine pathway (1). Host-derived signals increase cyclic GMP (cGMP) signaling in amphid neurons, which then secrete insulin-like peptides that stimulate synthesis of the steroid hormone dafachronic acid (DA). DA activates the nuclear receptor DAF-12 and promotes post-infection development (2, 3). Although volatile compounds from host skin and sweat attract L3i larvae (4, 5), the cues that trigger developmental activation after host entry remain poorly understood.

Olfactory receptors constitute the largest family of G protein-coupled receptors (GPCRs) and serve as chemical sensors in animals (6). Most vertebrate olfactory GPCRs are canonical odorant receptors (ORs), which belong to the rhodopsin-like class A family and are characterized by short N-terminal tails, conserved class A motifs, and ligand-binding pockets formed by their transmembrane domains (7). Nematodes have undergone an extensive expansion of a distinct family of chemosensory GPCRs that share structural features with vertebrate class A ORs (8, 9). These receptors detect diverse environmental cues, including food-associated chemicals and pheromones (10–13), but none has been shown to sense amino acids.

Here, we identify L-cystine as a host-derived signal that activates infective larvae of two parasitic nematodes whose hosts are evolutionarily distant: the human parasite *Strongyloides stercoralis*, which causes strongyloidiasis, a potentially life-threatening infection affecting more than 600 million people worldwide (14); and the insect parasite *Steinernema carpocapsae*. We further identify homologous class A odorant GPCRs in both species that respond to L-cystine. These findings reveal a conserved L-cystine-sensing system that allows parasitic nematodes to recognize the internal host environment and initiate development. They also suggest Ss-OR1 and related receptors as potential therapeutic targets for strongyloidiasis and other parasitic nematode infections.

## Results

### L-cystine activates *S. stercoralis* L3i larvae

Dulbecco’s Modified Eagle Medium (DMEM) activates feeding of *S. stercoralis* L3i larvae (15). To identify the active component, we used a feeding assay to test DMEM lacking amino acids, vitamins, salts, or NaHCO_3_. We separately tested the effect of omitting 5% CO_2_ from the incubation atmosphere. We added 25 mM HEPES to maintain pH under all conditions. Removing amino acids abolished L3i feeding, whereas removing the other components or omitting 5% CO_2_ had little effect (Fig. 1A). We therefore tested each of the 20 proteinogenic amino acids individually at 1 mM and found that only L-cystine strongly stimulated feeding (Fig. 1B).

**Fig. 1.**
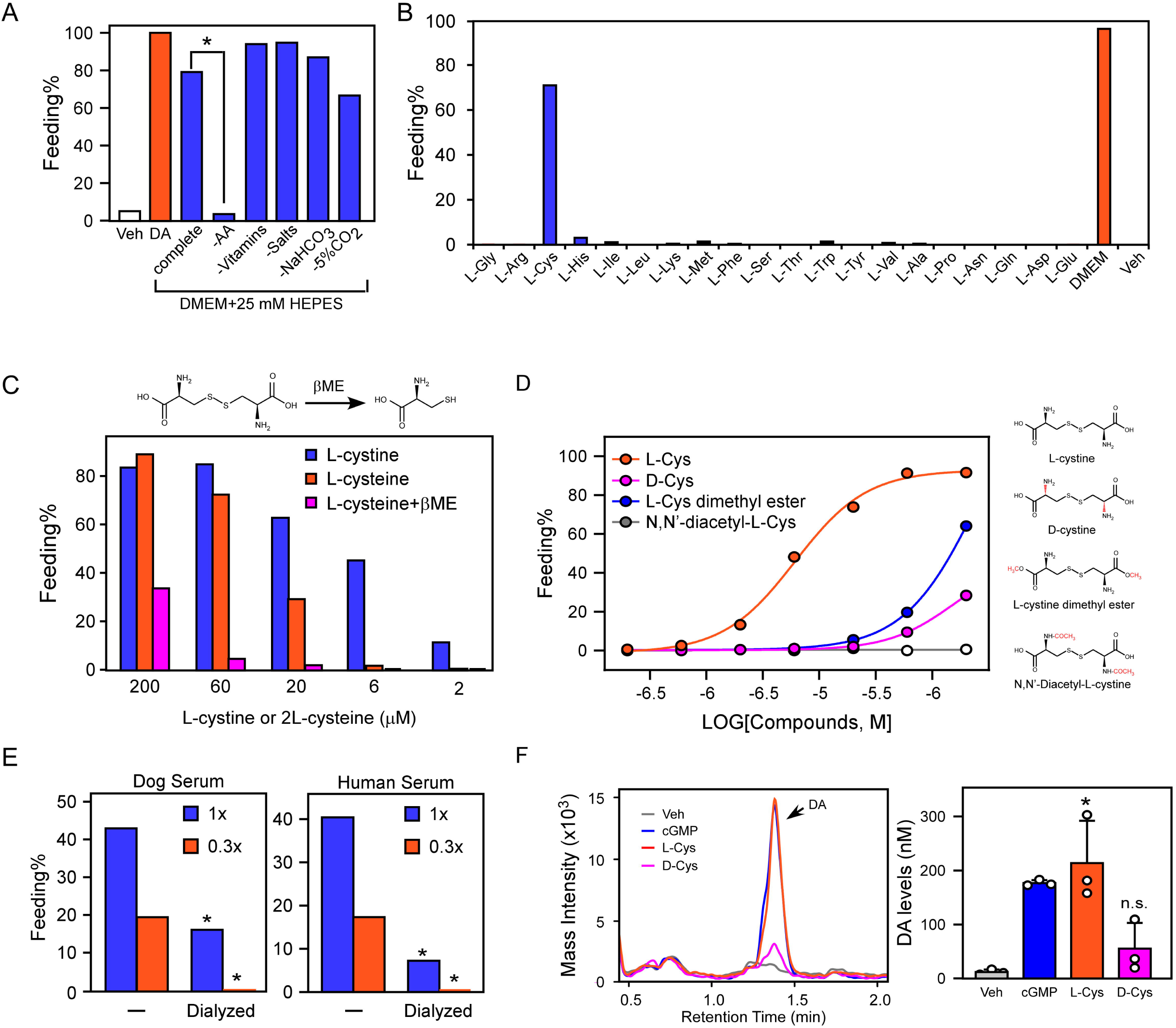
L-Cystine activates *S. stercoralis* L3i larvae. A, Analysis of DMEM components and CO_2_ for effects on L3i feeding. AA, amino acids; DA, Δ7-dafachronic acid. n = 60–100 larvae from two experiments. *, *P < 0.001* by Fisher’s exact test. B, L-cystine stimulates L3i feeding. All amino acids are at 1 mM. n = 170–280 larvae from three experiments. C, L-cystine, the oxidized form, activates L3i feeding more efficiently than L-cysteine. βME, β-mercaptoethanol. n = 50–200 larvae from two experiments. D, L-cystine stimulates L3i feeding in a concentration-dependent and structure-specific manner. n = 80–270 larvae from three experiments. Structural differences of L-cystine variants are highlighted in red. E, Activation of L3i feeding by host serum is reduced by dialysis. Sera were dialyzed against M9 buffer using a membrane with a 2-kDa molecular-weight cutoff. 1×, undiluted serum; 0.3×, 30% serum diluted in M9 buffer. n = 80–300 larvae from three experiments. *, *P < 0.001* compared with the corresponding undialyzed serum by Fisher’s exact test. F, L-cystine stimulates DA synthesis in *S. stercoralis* L3i larvae. n = 3; data are means ± SD; *, *P < 0.01*, n.s., not significant compared with the vehicle-treated group by Student’s t-test.

L-cystine consists of two L-cysteine molecules joined by a disulfide bond under oxidizing conditions. Because our assay conditions favor L-cystine, small amounts of L-cysteine could still be present and contribute to the response. We therefore compared L-cystine with L-cysteine and added β-mercaptoethanol (βME) to limit oxidation of L-cysteine. L-cysteine was less active than L-cystine, particularly in the presence of βME (Fig. 1C), supporting L-cystine as the more active form.

Having identified L-cystine as the active amino acid, we examined the concentration dependence and structural selectivity of the response. We compared L-cystine with D-cystine, L-cystine dimethyl ester, and N,N-diacetyl-L-cystine. The latter two compounds modify the carboxyl and amino groups of L-cystine, respectively. Only L-cystine strongly activated feeding, with an EC_50_ of approximately 6 µM (Fig. 1D). This value is below the reported circulating L-cystine concentration of 40–100 µM in human plasma (16–18), indicating that physiological concentrations are sufficient to activate the larvae. Both dog and human sera also activated L3i larvae, whereas dialysis markedly reduced this activity (Fig. 1E). These findings indicate that a small, dialyzable serum component contributes to larval activation and are consistent with circulating L-cystine serving as a host-derived signal.

Feeding is one measure of L3i activation, whereas activation also induces synthesis of dafachronic acid (DA), which activates the nuclear receptor DAF-12 and promotes reproductive development (2, 3, 19, 20). We therefore measured DA after treatment with L-cystine or D-cystine. L-cystine increased DA concentrations to approximately 200 nM, whereas D-cystine had a much smaller effect (Fig. 1F). The L-cystine response was similar to that produced by the membrane-permeable cGMP analog, 8-Br-cGMP. This concentration of DA occurs in developing parasites (2) and is sufficient to activate DAF-12 (3, 19, 20). Thus, L-cystine induces both feeding and DA synthesis in *S. stercoralis* L3i larvae.

### Identification of an L-cystine receptor in *S. stercoralis*

In *C. elegans*, GPCRs expressed in amphid neurons detect environmental signals and regulate development (1, 8). The structural selectivity of the L-cystine response suggested that a similar receptor-mediated mechanism operates in *S. stercoralis* (Fig. 1D). To identify the receptor, we first searched the genomes of *S. stercoralis* and several other nematodes using 31 human sensory or amino acid-responsive GPCRs as queries. This analysis identified 43 putative GPCRs in *S. stercoralis* and 179 in the other nematodes. We aligned and clustered all query and hit sequences (Table S1), then excluded 12 *S. stercoralis* GPCRs related to receptors with known functions, including glutamate and serotonin receptors. We ranked the remaining 31 candidates by their sequence identity to GPCRs in other nematodes (Table S2). Because no GPCR had previously been identified as an L-cystine receptor, we prioritized the least conserved candidates, reasoning that an unrecognized receptor might be evolutionarily divergent.

We used a CRISPR/Cas9 loss-of-function screen to test the five highest-priority candidates by inserting a GFP cassette into each gene through homology-directed repair ((21, 22); Fig. 2A). We scored feeding in GFP-positive F1 progeny exposed to DMEM or L-cystine and then genotyped the same larvae individually by single-worm PCR. Disruption of the fifth candidate tested, a putative class A GPCR that we designated Ss-OR1 (WormBase ID SSTP_0000730700; Fig. S1), impaired feeding in response to both DMEM (Table S2) and L-cystine (Fig. 2B). Feeding in response to L-cystine was reduced by approximately 60% among GFP-positive larvae in the Ss-OR1 CRISPR experiment (Fig. 2B). Approximately 75% of these larvae carried heterozygous or homozygous insertions at the *Ss-OR1* locus; the remaining GFP-positive larvae lacked a detectable insertion, consistent with GFP expression from nonintegrated repair-template DNA (Fig. 2A). Thus, Ss-OR1 is a major contributor to L-cystine-induced feeding initiation. Importantly, the cGMP analog 8-Br-cGMP restored feeding in the edited population, indicating that the larvae remained capable of feeding and that their downstream activation machinery was intact (Fig. 2B). By contrast, disruption of the closest paralog of Ss-OR1, Ss-OR3, or either of two unrelated GPCRs, Ss-GPR9 and Ss-GPR22, did not impair feeding (Table S2). Together, these results show that Ss-OR1 is required for a normal L-cystine response and are consistent with Ss-OR1 acting upstream of cGMP signaling.

**Fig. 2.**
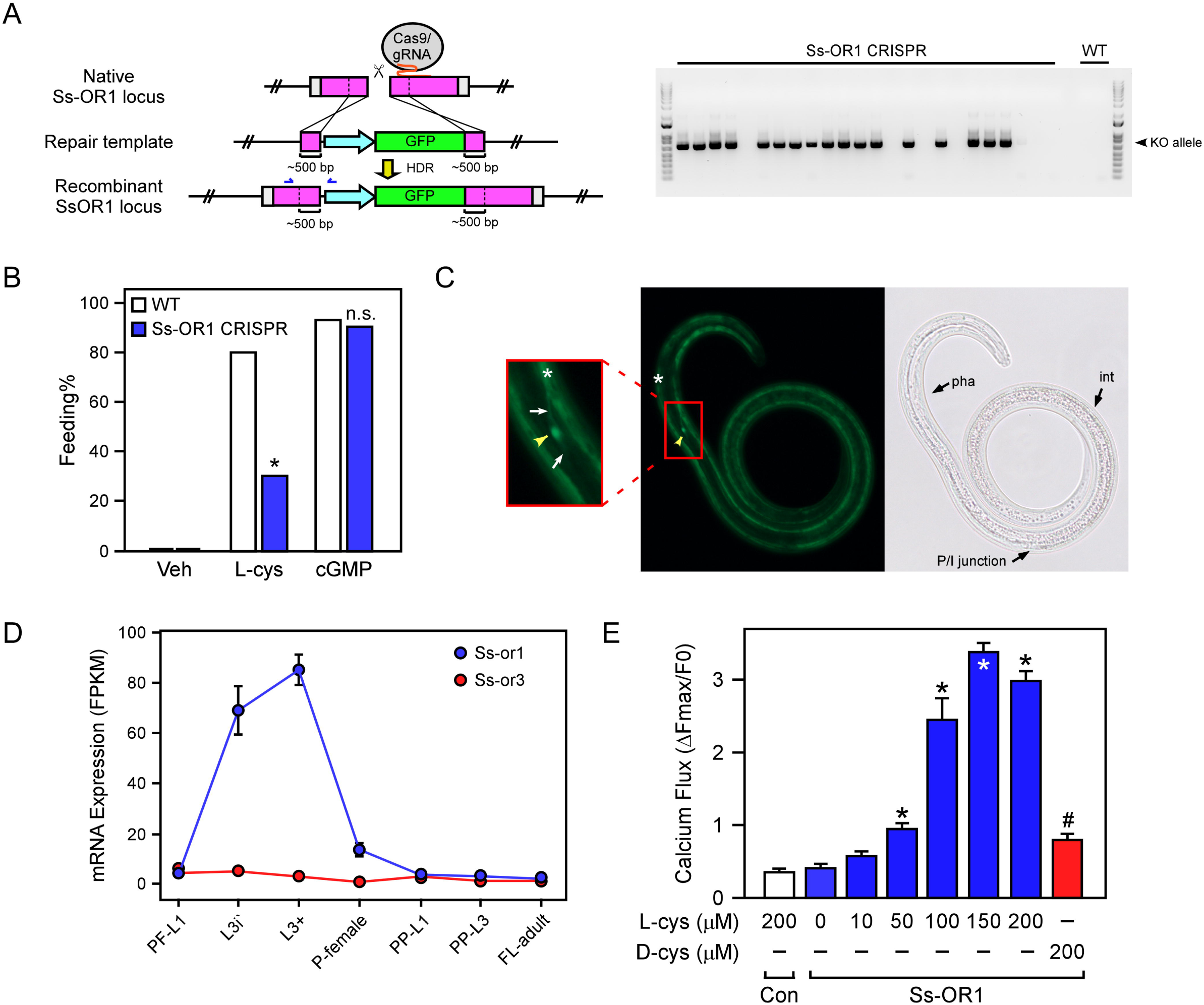
Ss-OR1 is an L-cystine receptor that mediates L3i activation in *S. stercoralis*. A, CRISPR-induced Ss-OR1 gene disruption. Plasmids expressing Cas9 and sgRNA, together with repair-template plasmids, were injected into free-living females. GFP-positive F1 L3i progeny were assayed for feeding. After feeding was scored, the same larvae were individually genotyped by single-worm PCR. Blue arrows indicate the primer sets used for genotyping. B, Ss-OR1 is required for normal L-cystine-induced L3i activation. Wild-type or Ss-OR1 CRISPR L3i larvae were treated as indicated, and feeding behavior was measured. n = 30–60 larvae from three experiments. *, *P* < 0.001, n.s., not significant compared with the corresponding wild type by Fisher’s exact test. C, Ss-OR1 is expressed in a presumptive amphid neuron in L3i larvae. Arrowhead, cell body; arrow, neurite; *, nonspecific autofluorescence; pha, pharynx; int, intestine; P/I junction, pharyngeal-intestinal junction. D, Ss-or1 mRNA expression is highest in the L3i and L3+ stages. Ss-OR3 is the closest homolog of Ss-OR1 in *S. stercoralis*. PF-L1, post-free-living L1; P-female, parasitic female; PP-L1/L3, post-parasitic L1/L3; FL-adult, free-living adult. E, Ss-OR1 is activated by L-cystine in a concentration-dependent manner in Sf9 cells. Control cells expressed human cytochrome P450 oxidoreductase. *, P < 0.001 compared with the vehicle-treated cells; #, P < 0.001 compared with the 200 µM L-cystine-treated cells, by Student’s t-test. Data are means ± SEM from 50–60 cells and are representative of three independent experiments with similar results.

### Ss-OR1 is expressed in a presumptive amphid sensory neuron

To determine where in the larvae Ss-OR1 may function, we examined its spatial expression pattern. We fused a 1.7-kb Ss-OR1 promoter fragment to GFP and generated transgenic L3i larvae. GFP was detected in a single cell near the pharyngeal bulbs and posterior to the nerve ring (Fig. 2C). Neurite-like processes extended from the cell body, whereas no comparable signal was observed in wild-type controls (Fig. S2).

This asymmetric expression pattern resembles that of many nematode chemoreceptors (9). Together, the location and morphology of the labeled cell are consistent with those of an amphid sensory neuron. In *C. elegans*, amphid neurons regulate exit from the L3i-like dauer stage (23).

We next asked whether Ss-OR1 expression is restricted to the stages that encounter the host. Ss-OR1 expression was highest in L3i and L3+ larvae, the latter representing L3 larvae activated within the host, and was low at other developmental stages ((24); Fig. 2D). The closest paralog, Ss-OR3, also remained low in L3i and L3+ larvae. Thus, both the spatial and temporal expression patterns of Ss-OR1 are consistent with a role in sensing the host environment.

### L-cystine activates Ss-OR1 in a heterologous system

Since the genetic and expression data implicated Ss-OR1 in L-cystine sensing, we tested whether L-cystine activates the receptor in a heterologous system. We expressed Ss-OR1 in *Spodoptera frugiperda* Sf9 cells, confirmed its cell-surface expression (Fig. S3), and measured cytosolic calcium during L-cystine treatment. L-cystine produced a concentration-dependent calcium signal in Ss-OR1-expressing cells, with an EC_50_ of approximately 90 µM (Fig. 2E, Movies S1–S3). In contrast, cells expressing the unrelated control protein human P450 oxidoreductase did not respond, and D-cystine produced a much smaller response in Ss-OR1-expressing cells. These results show that Ss-OR1 expression confers a selective cellular response to L-cystine.

### L-cystine responsiveness extends to an insect-parasitic nematode

We next asked whether this sensory mechanism extends beyond *Strongyloides*. Homologs of Ss-OR1 occur in several parasitic nematode lineages, including not only *Strongyloides* and *Parastrongyloides* species but also *Steinernema* species, hookworms, and other nematodes (Fig. 3A). We focused on *Steinernema* because it belongs to nematode clade IV with *S. stercoralis* (25) and infects insects through an L3i-like stage called the infective juvenile (IJ) (26, 27).

**Fig. 3.**
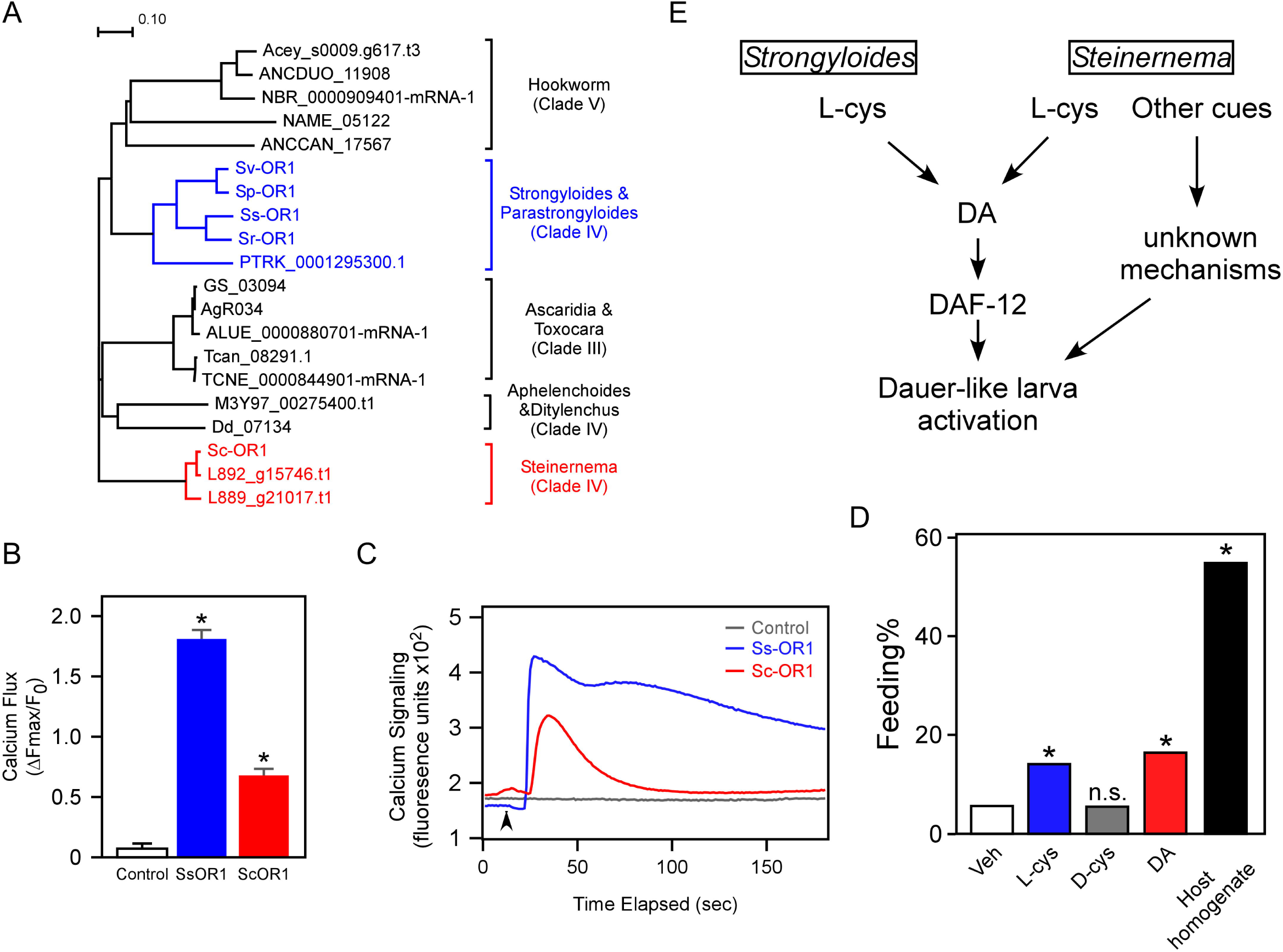
Conservation of the L-cystine/GPCR sensory pathway in parasitic nematodes. A, Neighbor-joining phylogenetic tree of Ss-OR1 homologs in parasitic nematodes. B, Ss-OR1 and Sc-OR1 are activated by L-cystine. Cells expressing Ss-OR1 or Sc-OR1 and control cells expressing human cytochrome P450 oxidoreductase were treated with 200 µM L-cystine. *P < 0.001 compared with control cells by Student’s t-test. Data are presented as means ± SEM from 50–60 cells and are representative of three independent experiments with similar results. C, Representative real-time calcium traces from control, Ss-OR1-expressing, and Sc-OR1-expressing Sf9 cells. The arrowhead indicates addition of 200 µM L-cystine. D, L-cystine partially activates infective juveniles of *S. carpocapsae*. Infective juveniles were treated with vehicle, L-cystine, D-cystine, DA, or host homogenate, and feeding was measured. n = 420–500 juveniles from three experiments. *, *P < 0.001* compared with the vehicle-treated group by Fisher’s exact test. E, Proposed pathways governing the activation of developmentally arrested larvae in parasitic nematodes.

We tested Sc-OR1, an Ss-OR1 homolog from *S. carpocapsae* that shares approximately 32% sequence identity with Ss-OR1 (Table S4) and found that L-cystine activated it in Sf9 cells (Figs. S3 and 3B–C and Movie S4). L-cystine also stimulated feeding in developmentally arrested *S. carpocapsae* IJs (Fig. 3D). DA produced a similar response, consistent with activation of Sc-DAF-12 (Fig. S4). However, L-cystine and DA each produced only about one-third of the response to host homogenate (Fig. 3D), indicating that other host signals contribute to full larval activation. Thus, *S. carpocapsae* has both an L-cystine response and an L-cystine-responsive OR1 homolog. Together, these findings indicate that L-cystine sensing extends across nematode parasites with evolutionarily distant hosts.

## Discussion

Parasitic nematodes use amphid neurons to detect their environment. Heat, sweat odorants, sodium, and CO_2_ guide host-seeking behavior (4, 5, 22, 28), but much less is known about the signals that activate development after host entry. Known activators differ among species: temperature activates *Nippostrongylus brasiliensis*, whereas serum albumin and glutathione derivatives activate *Ancylostoma* species (5, 28–30).

Changes in proton concentration may also contribute to *S. stercoralis* activation: removing CO_2_ inhibits activation (5), but not when pH is held constant (Fig. 1A). Here, we identify L-cystine as a previously unrecognized host-derived signal that stimulates feeding by *S. stercoralis* L3i larvae, marking their transition from a developmentally arrested, environmentally motile stage to a parasitic stage adapted for growth within the host. We further identify Ss-OR1 as an L-cystine-responsive receptor that contributes to this developmental response. Several observations support the physiological relevance of this signaling pathway. L-cystine circulates in human plasma at 40–100 μM (16–18), overlapping the concentrations that activate Ss-OR1. It could therefore be encountered during or shortly after skin penetration, when L3i larvae begin to activate (15). Moreover, *Ss-or1* expression is highest in L3i and *in vivo* activated L3+ larvae (Fig. 2D) (24), the developmental stages positioned to detect and respond to host-entry signals. Thus, L-cystine–Ss-OR1 signaling provides a potential molecular link between entry into the mammalian host and initiation of the parasitic life cycle (Fig. 3E).

The ligand–receptor relationship extends to *S. carpocapsae*: L-cystine stimulated infective juvenile feeding, and the *S. carpocapsae* OR1 homolog responded to L-cystine. Thus, both L-cystine responsiveness and an L-cystine-responsive OR1 homolog are conserved. The weaker responses of *S. carpocapsae* IJs to L-cystine and DA than to host homogenate suggest that full activation requires additional host-derived cues, potentially including other components of insect hemolymph. This contrasts with the more robust response of *S. stercoralis* L3i larvae to L-cystine alone and may reflect adaptation to distinct host environments. This conservation is striking because Ss-OR1 and Sc-OR1 share only approximately 32% sequence identity, despite both species belonging to clade IV (25). BLAST analysis identified OR1 homologs with even lower sequence identity in the clade IV nematodes *Aphelenchoides* and *Ditylenchus* (29% and 28%, respectively; Table S4). Key residues within the ligand-binding pocket may therefore remain conserved despite low overall sequence similarity. Identifying these residues may help uncover additional amino-acid-sensing GPCRs in parasitic nematodes.

The identification of Ss-OR1 also reveals an unexpected mechanism for amino acid sensing. Most GPCRs known to detect amino acids belong to class C and function in processes such as neurotransmission and taste (31). In fish, class C GPCRs can function as olfactory receptors to sense a broad range of L-amino acids in the environment. For example, goldfish OR5.24 responds to 19 of the 20 proteinogenic amino acids (32, 33). The class A GPCRs GPR139 and GPR142 also respond to amino acids (34, 35), although the physiological relevance of amino acid sensing by these receptors remains uncertain (34–36). Ss-OR1 is distinct from these previously characterized receptors: it is a class A GPCR expressed in a presumptive sensory neuron and responds selectively to the oxidized amino acid L-cystine. Moreover, Ss-OR1 responds to L-cystine at concentrations near those found in host circulation (16–18), further supporting its function as a sensor of the internal host environment. Thus, Ss-OR1 defines a previously unrecognized mode of GPCR-mediated cystine sensing that parasites use to control a life-cycle transition.

We previously showed that DAF-12 controls *S. stercoralis* development and can be targeted pharmacologically (2). The present study extends this developmental pathway upstream by identifying Ss-OR1 as part of the sensory mechanism that initiates parasite activation after host entry. Disrupting *Ss-or1* impairs L-cystine-induced feeding, whereas cGMP bypasses this defect, placing Ss-OR1 upstream of a signaling pathway previously implicated in L3i activation (15). Because this transition is essential for establishment of infection, Ss-OR1 may provide a new opportunity to interfere with parasitism at its outset. An antagonist could prevent larvae from responding appropriately after host entry, whereas an agonist could prematurely trigger the developmental program outside the host. These complementary strategies raise the possibility of targeting the sensory decision that commits an infective larva to parasitic development. More broadly, our findings establish an oxidized amino acid as a host signal that triggers developmental activation in parasites of insects and humans.

## Materials and Methods

### *S. stercoralis* maintenance and preparation

The *S. stercoralis* UPD strain was maintained in dogs as previously described (38). The infection was established by subcutaneous injection of 5,000 L3i larvae into dogs treated daily with 0.25–1 mg/kg prednisolone. Feces of the dogs were then collected daily for fecal culture. L3i larvae of *S. stercoralis* were prepared as described (2, 37).

Briefly, feces from dogs were mixed with three volumes of charcoal and placed on dampened Whatman filter paper in petri dishes. The cultures were maintained at 25°C for 7 days, after which L3i larvae were collected using Baermann funnels. L3i larvae were then sterilized by washing 5 times with M9 buffer supplemented with 100 U/ml penicillin, 100 µg/ml streptomycin, and 50 μg/ml gentamicin. The larvae were then resuspended in M9 buffer for subsequent experiments.

### Bioinformatic analysis

Sequences in this study were extracted from Wormbase ParaSite and NCBI. Multiple sequence alignment was performed using Clustal Omega with default parameters. An evolutionary distance matrix was calculated, and a phylogenetic tree was constructed using the neighbor-joining method.

### Feeding assays

*S. stercoralis* L3i feeding assays were performed as previously reported (2, 38). Customized DMEMs were formulated based on recipes from Thermo Fisher (Waltham, MA; cat# A14430), and the components were purchased from Sigma Aldrich (St. Louis, MO) and Alfa Aesar (Ward Hill, MA). Unless otherwise stated, amino-acid-free custom DMEM supplemented with 25 mM HEPES (pH 7.3) and 100 U/ml penicillin-streptomycin (Thermo Fisher, Waltham, MA) was used as the vehicle for all treatments. Briefly, approximately 50-100 *S. stercoralis* L3i larvae were incubated with the indicated treatments at 37°C with 5% CO_2_ for 21 h. To visualize feeding behavior, treated larvae were incubated with 200 µg/ml Alexa Fluor 594 carboxylic acid (Thermo Fisher, Waltham, MA) at 37°C for 3 h, then washed five times with PBS and examined by fluorescence microscopy. Unless otherwise stated, 200 µM cystine and 10 µM Δ7-DA were used.

*S. carpocapsae* infective juveniles (IJs) were prepared and activated using the waxworm homogenate method described by Lu et al. (39) and modified for large-scale activation by Mishra et al. (40). To prepare the homogenate, frozen *Galleria mellonella* larvae were ground in liquid nitrogen, suspended in PBS and boiled three to four times with intermittent stirring. The supernatant was collected and supplemented with 1× penicillin-streptomycin-neomycin immediately before use. For larval feeding, ∼10,000 IJs were washed at least three times with sterile 0.8% NaCl containing 0.01% Triton X-100, then transferred to autoclaved sponge pieces soaked in waxworm homogenate or in PBS containing vehicle, 1 mM L-cystine, 1 mM D-cystine, or 10 µM Δ7-DA. To visualize the feeding, 100-200 of the treated IJs were incubated with 200 µg/ml Alexa Fluor 594 carboxylic acid (Thermo Fisher, Waltham, MA) at 16°C for 3 h, then washed five times with PBS and examined by fluorescence microscopy.

Data from independent experiments were pooled and presented as the percentage of larvae feeding, and statistical significance was tested by Fisher’s exact test.

### Quantitative analysis of DA concentrations

DA measurement was performed as previously described (2). Briefly, approximately 10,000 L3i larvae were incubated with the indicated treatments at 37°C with 5% CO_2_ for 3 days. For single-point quantification, 100 nM Δ7-DA was added to untreated L3i larvae as an internal standard. Lipids were then extracted from the larvae using the Folch extraction method and dried under a nitrogen stream. Δ7-DA was derivatized to Δ7-DA-picolylamine (Δ7-DA-PA) for analysis by incubating lipids with 3.3 mM triphenylphosphine, 3.3 mM 2,2′-dipyridyl disulfide, and 333 ng/µl 2-picolylamine at 60°C. The samples were then loaded onto a C18 column (Phenomenex, Torrance, CA) and eluted with a gradient of acetonitrile (40–60%) in water using UPLC (Shimadzu, Columbia, MD). Δ7-DA-PA was detected using a SCIEX Triple Quad 4500 mass spectrometer in positive multiple reaction monitoring (MRM) mode with m/z transition 505 → 487 and quantified using the software MultiQuant (AB Sciex LLC, Framingham, MA) by comparing Δ7-DA-PA peak areas. Data from three experiments are presented as means ± SD with statistical significance determined by Student’s t-test. Unless otherwise stated, 0.5 mM 8-Br-cGMP, 200 µM L- or D-cystine and 10 µM Δ7-DA were used.

### Ss-or1 expression in L3i larvae

mRNA expression of Ss-or1 and Ss-or3 was obtained from the JBrowse tool implemented in parasite.wormbase.org, as first reported (24). For Ss-OR1 reporter expression, the 1,685-bp upstream fragment from the start codon of Ss-or1 was amplified and cloned into the pAJ09 plasmid using HindIII and XmaI restriction sites to replace its original Ss-gpa3 promoter, generating a GFP reporter driven by the Ss-or1 promoter. The construct was then microinjected with the co-injection marker pAJ50-2 (*Sr-eft-3p:mRFP*) into the gonads of free-living females, which were then cultured with male adults in feces at 25°C for 7 days. The resulting L3i larvae were collected by Baermann funnels, mounted on 2% agarose gel pads, and imaged using a fluorescence microscope (Zeiss Imager A2).

### CRISPR-mediated gene knockout in L3i larvae

To disrupt expression of the *S. stercoralis* genes, CRISPR-mediated gene editing was performed using homology-directed repair (HDR) as outlined (Fig. 2A) and previously reported (2, 5, 38). Briefly, single-guide RNAs (sgRNAs) targeting the indicated genes were designed by SSC (http://crispr.dfci.harvard.edu/SSC/) or CHOPCHOP (http://chopchop.cbu.uib.no/), and potential off-target effects were analyzed by BLAST against the *S. stercoralis* genome (summarized in Table S3). The sgRNAs were then cloned into the gRNA vector pMLC60 (5) to obtain gRNA plasmids. For each target gene, the homology-directed repair template plasmid was made by cloning ∼500 bp homology arms flanking the Cas9 cutting sites into pPV529 vector, which carries a GFP-expressing cassette (*Sr-eef-1p::GFP::era3’UTR*).

Approximately 50 free-living females of *S. stercoralis* were microinjected with plasmids expressing gRNA (60 ng/µl), Cas9 (pPV540, 20 ng/µl), and the corresponding homology-directed repair template plasmids (10 ng/µl). Injected females were then placed with adult males in the fecal culture and the F1 progeny from these matings were collected after a 7-day fecal culture at 25°C using the Baermann technique. GFP-positive F1 L3i larvae were manually picked and assayed for feeding. After feeding was scored, the same larvae were recovered and individually genotyped by single-worm PCR using oligonucleotides that annealed to the 5′ regions outside of the homology arm and to the GFP-expressing cassette (Table S3). Data from at least three experiments were pooled together and are presented as the percentage of feeding, with statistical significance tested by Fisher’s exact test.

### Calcium flux assays

For expression, Sf9 codon-optimized Ss-OR1 and Sc-OR1 were fused to an N-terminal HA tag and cloned into pFastBac Dual vectors containing sequences encoding GP64 signal peptides. Baculoviruses expressing HA-SsOR1 or HA-ScOR1 were then prepared according to the manufacturer’s instructions (Thermo Fisher, Waltham, MA). For calcium flux assays, Sf9 cells were cultured in 35-mm glass-bottom dishes (Cellvis, Mountain View, CA) and were infected with baculovirus carrying the indicated genes for 24 h. After a 2 h starvation in HBSS (Thermo Fisher, Waltham, MA), cells were subsequently loaded with 2 µM calcium indicator Fluo-4 AM (Thermo Fisher, Waltham, MA) in ECB buffer (20 mM HEPES, pH 7.4, 125 mM NaCl, 5 mM KCl, 1.5 mM MgCl_2_, 1.5 mM CaCl_2_, 10 mM glucose, 2.5 mM probenecid) for 20 min. Following ECB buffer washing and 30 min incubation in the dark, the cells were live-imaged for calcium signals every 1 s using a Nikon CSU-W1 fluorescence confocal microscope for 3 min. The indicated treatments were administered 10-12 seconds after the imaging began.

The images were converted to movies at 6 frames/s with Image J. Unless otherwise stated, 200 µM L- or D-cystine was used.

For data analysis, fluorescence intensities were quantified in 50–60 cells at the initial time point (F0) and at each subsequent time point [F(t)]. The normalized fluorescence change at each time point was calculated as [F(t) − F0]/F0. The maximum normalized change for each cell is presented as the mean and SEM. Significance was assessed using Student’s *t* test. The experiment was repeated at least three times with similar results.

### Statistical analysis

Statistical analyses were conducted as described (2, 3) using GraphPad QuickCalcs for binomial data (Fisher’s exact test) and Microsoft Excel for continuous data (two-tailed Student’s t-test with equal variance). P values less than 0.01 were considered statistically significant. Dose-response curves were plotted using SigmaPlot 11.0 and the curves were fitted to a four-parameter sigmoidal equation.

## Acknowledgments

We thank Xuewu Zhang for discussions. This work was supported by the NIH (R35GM137934 to A.R.D. and R01AI167967 to J.J.C.), the Robert A. Welch Foundation (I-1948-20240404 to J.J.C., I-1275 to D.J.M.), Lyda Hill Philanthropies (Hill Prize to S.A.K. and D.J.M.) and the Howard Hughes Medical Institute (J.J.C. and D.J.M.).

